# Cellular And Molecular Effects Of Understudied Kinase Pregnancy-Upregulated Non-Ubiquitous Calcium-Calmodulin Dependent Kinase (PNCK) In Renal Cell Carcinoma

**DOI:** 10.1101/2022.05.09.491212

**Authors:** Derek J. Essegian, Valery Chavez, Floritza Bustamante, Stephan C. Schürer, Jaime R. Merchan

**Author notes:** Equal contribution (Derek J. Essegian, Valery Chavez).

## Abstract

Renal Cell Carcinoma (RCC) is a uniformly fatal disease when advanced. While immunotherapy and tyrosine kinase inhibitor-based combinations are associated with improved progression-free and overall survival, the majority of patients eventually develop treatment resistance and succumb to progressive, refractory disease. This underscores the urgent need to identify novel, non-canonical RCC targets for drug development. Through a comprehensive pan-cancer, pan-kinome analysis of the Cancer Genome Atlas (TCGA), the understudied kinase, pregnancy upregulated non-ubiquitous calcium-calmodulin dependent kinase (PNCK) was identified as the most differentially overexpressed kinase in RCC. PNCK mRNA was significantly overexpressed in RCC tissues compared to adjacent normal tissue, and its overexpression correlated with tumor T-stage grade and poor disease specific survival in both clear cell and papillary RCCs. PNCK overexpression in VHL mutant and VHL wild type RCC cell lines was associated with increased CREB phosphorylation, as well as increased cell proliferation and cell cycle progression. PNCK down-regulation, conversely, was associated with inhibition of CREB phosphorylation, decreased cell proliferation, cell cycle arrest and increased apoptosis, with differential effects observed between VHL mutant and VHL wild type cell lines. Pathway analyses in PNCK knockdown cells showed significant down regulation of hypoxia and angiogenesis pathways, as well as modulation of pathways promoting cell cycle arrest and apoptosis. The above results demonstrate for the first time the biological role of PNCK, an understudied kinase, in renal cell carcinoma and validate PNCK as a potential novel target for drug development in this fatal disease.

## INTRODUCTION

Renal Cell Carcinoma (RCC) is an aggressive malignancy expected to affect 79,000 new Americans in 2022 of which an estimated 13,920 are expected to succumb to this disease[1]. Advances in the understanding of RCC pathogenesis [2], including VHL silencing, HIF accumulation, increased angiogenesis and the RCC immune microenvironment [3], have led to the development of new treatment options, including Tyrosine Kinase Inhibitors (TKIs), mTOR inhibitors and more recently, immune checkpoint inhibitors (CPIs), given alone or in combination [4]. While these agents have significantly improved progression free and overall survival in advanced clear cell RCC, a large number of patients eventually develop treatment resistance, leaving patients with few treatment options and a poor prognosis [5]. Therefore, identifying novel targets for RCC drug development is an urgent, unmet need.

The calcium-calmodulin kinase (CAMK) family as a whole is understudied with currently no approved drugs or clinical trials targeting these kinases[6, 7]. PNCK (CAMK1B), an isoform of CAMK1, is among the least characterized CAMK members, despite emerging studies demonstrating its prognostic and potential therapeutic relevance [8–12]. PNCK is a 38-kDa protein whose catalytic domain shares 45-70% sequence identity with other members of the multifunctional CAMK family [13]. PNCK is temporally expressed and restricted to very few tissues through development and into adulthood, with highest expression in the dentate gyrus in the hippocampus[14, 15]. The *in vivo* substrates of PNCK are unknown. However, *in vitro* studies suggest that PNCK phosphorylates CAMK1A, CREB, ATF1 and Synapsin [16]. PNCK has been linked to aggressive phenotypes and poor outcomes in limited studies in breast (BC), renal cell (RCC), nasopharyngeal, and hepatocellular carcinomas [8–12]. In clear cell RCC, PNCK expression (by immunohistochemistry) was shown to be significantly higher in tumor compared to adjacent normal tissues and was established as an independent predictor of poor survival, tumor size and histological grade [9]. Studies in breast cancer suggest that PNCK stimulates proliferation, clonogenicity, cell cycle progression and treatment resistance [17], and that it may play a role in modulating the tumor microenvironment [8]. To date, however, there have been no comprehensive studies on PNCK, CAMKs and the CAMK pathway in renal cell carcinoma.

Towards our goal to find new RCC targets, we performed an integrative analysis of available cancer databases (TCGA and Pharos) to identify understudied kinases that are overexpressed in RCC[7]. Our analysis identified Pregnancy-Upregulated Non-ubiquitous Calcium/Calmodulin dependent kinase (PNCK), as the most differentially overexpressed kinase in RCC, and was significantly associated with poor clinical outcomes. The current report aims to characterize the *in vitro* biological effects and mechanisms of PNCK up and down-regulation in renal cell carcinoma models. Our results suggest that PNCK is a promising target for novel drug development efforts in this fatal disease.

## MATERIALS AND METHODS

### TCGA Transcriptomics Analysis

Analysis of Pharos and TCGA data was executed as described in Essegian et. al[7]. Understudied kinases were identified in Pharos and were queried in TCGA for differential expression. Transcriptomic data was normalized using R package Limma and differential expression was calculated using TCGAbiolinks R package [18] with RSEM converted to Log2 expression. Only primary tumor samples were included in the analyses-excluding secondary or metastatic tumor samples. KIPAN (Pan-Kidney) analyses were done combining KICH, KIRC and KIRP datasets. Kaplan-Meier (KM) curves were generated using Prism and Xena[19] . Cohorts for KM analyses were split into high and low expressers based on the median expression PNCK. Statistical analyses were calculated using Prism-including students t-test and ANOVA to determine difference in expression between normal and tumor tissues and amongst tumor grades and stages.

### Cell Culture

A-704 (ATCC® HTB-45™), 786-O (ATCC® CRL-1932™), ACHN (ATCC® CRL-1611™), Caki-1 (ATCC® HTB-46™), HEK293 (ATCC CRL-1573TM), A498 (ATCC® HTB-44™), HREC (ATCC® PCS-400-012) and HUVEC (ATCC® CRL-1730™) cells were all obtained from ATCC. ACHN, A704 and HEK293 cells were cultured using EMEM (Eagle’s minimal essential medium). HUVEC cells were grown with F-12K Medium with Heparin and Endothelial cell growth supplement. 786-O and A498 cells were cultured with RPMI 1640 Media. Caki-1 cells were cultured with McCoy’s 5A Modified Medium. All cell media was supplemented with 10% Fetal Bovine Serum and 1X penicillin/streptomycin. For all transfections, antibiotics were excluded from media. Cells were grown and maintained at 37°C, 5% CO2. Frozen cultures were stored at -80C or long-term in liquid nitrogen.

### Real-time quantitative PCR and analysis

Total RNA was extracted from cells grown to 80% confluence using the Qiagen (Hilden, Germany) RNAeasy Mini Kit (Cat No: 74104). Quantity and purity of RNA was assessed using the ThermoFisher Scientific (Waltham, MA, USA) NanoDrop OneC Microvolume UV-Vis Spectrophotometer. RNA was reverse transcribed into cDNA using Qiagen Quantitect Reverse Transcriptase kit (Cat No: 205311). cDNA was diluted to 50ng/uL and stored at -20C until further use. qPCR primers were designed from IDT (Coralville, IA, USA) against ref seq NM_001039582(2) which targets exons 9-11 and detects all variants and isoforms of PNCK. Forward: TTTGACTCT CCTTTCTGGGATG. Reverse: GTT GGCAGGTGAACCTCTT. cDNA was amplified using 10ng of cDNA and IDT PrimeTime Gene expression Master Mix (#1055772) and a final reaction volume of 20uL. qPCR experiments were run on the Biorad CFX96 and gene expressions were measured relative to GAPDH. No reverse transcriptase (NRT) and no template control (NTC) were used to evaluate the signal contamination of genomic DNA or background from primers. The amplification conditions for the quantitative PCR (qPCR) reactions were as follows: 1 cycle of initial denaturation at 95°C for 3 min followed by 40 cycles of denaturation at 95°C for 30 s, annealing at 60°C for 1min. All reactions were performed in quadruplicate. qPCR data was analyzed by the comparative threshold (ΔΔ*C_t_*) method. The derived ΔΔ*C_t_* values were converted into fold-difference values.

### SDS protein gel and Western Blot analysis

Protein concentration was quantified using the Bradford protein assay (#23236, ThermoFisher Scientific, (Waltham, MA, USA) following manufacturer’s recommendations. Western blots were run using 25-50ug of protein per well with pre-cast gradient gels. Gels were treated with 8M Urea and transferred to nitrocellulose film overnight at RT. Membranes were blocked for two hours with 5% BSA or 5%/ milk (depending on manufacturers suggestions for antibody). For PNCK (# ab235093, Abcam Cambridge, MA, USA), membranes were blocked with BSA and incubated overnight with primary antibody at a 1:500 dilution. Membranes were then washed 3x with TBS/Tween and incubated at RT for 1h with secondary antibody. Chemiluminescence peroxidase activity was revealed with theThermo Scientific SuperSignal West kit. Bands were visualized using ChemiDoc™ MP Imaging System (Biorad, Hercules, CA, USA). Protein loading was determined by GAPDH analysis (# ab181602, 1:10000 dilution; Abcam Cambridge, MA, USA).

### Generation of PNCK overexpressing cell lines

Stable PNCK overexpressing cells were generated via lentiviral transduction Lenti ORF particles of PNCK (transcript variant 1) were obtained from Origene (Rockville, MD, USA) (Cat #RC229215L4) at a titer of >1 x10^7 TU/ml. PNCK ORF and control particles were built with the pLenti-P2A-Puro vector. Cells were plated in a 48-well plate to 50% confluency and were infected with the particles at an MOI of 25 as per manufacturers recommendations. At 48h post-infection, GFP positivity was detected via fluorescence microscopy. Cells were then split and single colonies were isolated and selected with puromycin (1-2ug/ml) for 7-10 days. After first round of single colony selection, cells were sorted using flow cytometry sorting for Top1% of GFP expression and then a second round of single colony selection (from to 1% selected cells PNCK Overexpression was confirmed via qPCR and western blot and colonies with the highest level of expression were chosen for further experiments.

### PNCK knock-down

Double stranded siRNAs (dsiRNA) were obtained from IDT DNA Technologies (Coralville, IA, USA). The predesigned sequences were from the TriFECTa Kit which consisted of 3 PNCK dsiRNAs (13.1 rArArG rArUrC rArUrG rGrUrC rUrCrU rGrArC rUrUrU rGrGA C, 13.2 rArCrA rUrCrA rGrCrA rGrCrG rUrCrU rArCrG rArGrA rUrCC G, 13.3, rCrGrG rArUrC rUrCrG rUrArG rArCrG rCrUrG rCrUrG rArUrG rUrCrC), Negative controls and resuspension buffer. To determine the most potent and selective dsiRNA, all three were tested (13.1, 13.2 and 13.3) at concentrations from 1 nM to 20 nM. Cells were grown in 10cm plates to 60-70% confluence and dsiRNA was added using a mixture of Optimem Serum-Free Media and Lipofectamine RNAiMax (#13778030, ThermoFisher Scientific, Waltham, MA, USA). At 24h, Cy3-labeled scrambled transfection controls were used to confirm >90% transfection efficiency via microscopy. PNCK knock down was confirmed at different time-points post-transfection via qPCR, western blot and immunofluorescence. Dose response curves determined that siRNA #2 delivered the greatest levels of knockdown with low toxicity at 10nm. For A498 cells, the greatest knockdown was achieved using siRNA #1 at 20nM. Increasing the concentration >25nM to 50nM did not result in increased levels of knockdown.

### Immunofluorescence (IF)

40,000 cells/well were seeded in Labtek slides, followed by fixation in Formalin after overnight incubation. After fixing, the background produced from formalin was quenched with NH4Cl for 15min and then samples were blocked and permeabilized with 0.5% Goat serum + Triton-X 0.3% in PBS for 2h, prior to overnight incubation with primary antibodies: (p-CREB: 1:1000 (Cell Signaling 9198), PNCK 1:250 (Invitrogen PA5-99601) Angiopoietin 1 (Abcam 133425) 2.5µ/ml, Angiopoietin 2 (Abcam 153934) 1:500, Cyclin D1 (Abcam 16663) 1:150, p21 (Abcam 109520) 1:100, p27 (Cell Signaling 3886) 1:800, Cyclin A (Cell Signaling 67955) 1:500, Cyclin B2 (Abcam 185622) 1:300, Ki67 (Abcam 15580) 0.5µ/ml.Alexa Fluor 555 conjugated secondary antibodies (1:350) were used to visualized cells under fluorescent microscopy for overexpression experiments. Alexa Fluor 488 was used in dsiRNA knockdown experiments.

### XCelligence Real Time Cell Analysis (RTCA) Cell Proliferation assay

Cells were grown in T75 flasks to 60-80% confluency. The xCelligence RTCA (Agilent and ACEA, Santa Clara, CA, USA) 96- well electronic microtiter plate (E-plate) was first “blanked” with 100uL of cell culture media in every well to test background electrical impedance. Cells were then trypsinized and counted using a hemocytometer to plate 6,000-10,000 cells per well in a 96-well xCelligence E-plate in an additional 100ul of culture media (for a final well volume of 200uL). Growth was monitored in real-time for up to 120 hours with impedance assessed at 15-30-minute intervals using the xCelligence Real-time Cell Analyzer (RTCA). Data was exported from the xCelligence system and analyzed per individual well. Cell growth results were shown as bar graphs showing the averages (+/- SD) of experiments done at least in sextuplicate.

### Cell Cycle Analysis by Flow Cytometry

To assess the effect of PNCK activity on cell cycle progression, flow cytometry was utilized to quantify the relative populations of cells in each stage of the cell cycle. For overexpression experiments, Cells were grown in 10cm plates to 60-80% confluence and were trypsinized and washed 3x with ice cold PBS. Cells were then fixed and permeabilized with 70% ethanol overnight at -20C. For knockdown cells, cells were transfected first with dsiRNA and then after desired time points post-transfection, cells were then fixed and permeabilized with 70% ethanol overnight at -20C. Samples were then treated with a mixture of RNAse-H and Propodium Iodine and incubated at 37°C for 1 hour. Using the LSR-II Cytometer, samples were analyzed. 50-100K events were recorded at a low-voltage setting with a slow flow rate. Side scatter and forward scatter gating was employed to eliminate both apoptotic cells and clumps of cells so that only single cells were used in the final analysis. FCS files were exported and analyzed using FlowJo v.10.6.2 (https://www.flowjo.com/solutions/flowjo). Experiments were run in sextuplicate and repeated at least 3 times. Student’s T-test was used to determine significant changes in cell cycle population ratios in treated and control cells.

### RT^2^ Profiler Cancer PathwayFinder Array

Cell lines were treated with siRNA or control siRNA as describes above with 3 biological replicates and 3 technical replicates. At 72h post transfection, RNA was extracted and purified as described above. RNA was reverse transcribed using the RT2 First Strand Kit (Qiagen, 330401), and cDNA was then added with SYBR green mastermix (Qiagen, 330504) to 96- well RT2 Profiler Human Cancer PathwayFinder PCR Array plates (Qiagen, PAHS-033Z), according to the manufacturer’s instructions. Quantitative PCR was run as described previously and data was analyzed using Qiagen’s GeneGlobe Data Analysis center, using both ACTB and GAPDH as housekeeping genes. Input genes for each cell line studied were either significantly up-regulated or downregulated genes from the RT2 Profiler Cancer PathwayFinder array with a p-value <0.05.

### Caspase Glo 3/7 Apoptosis Assay

Cell lines were treated with siRNA or Cy3-control siRNA at 10nM as described above in replicates of 6 and were transferred to 96-well plates. For A498 cells, siRNA #1 was used at a concentration of 20nM. Cell count was normalized to 8,000 cells per well prior to adding Glo reagent at 72h post transfection, caspase activity was detected using Caspase Glo 3/7 Apoptosis Assay and Promega Luminometer. Luminescence signal was corrected with blank and untreated cells.

### Statistical Analysis

In vitro data are presented as meansL±LSD. Results from in vivo studies are shown as meansL±LSEM. All in vitro experiments were performed at least in triplicate unless otherwise specified and repeated at least twice. Statistical analysis among groups was performed by ANOVA, and sub-group comparisons were made with the student’s T-test, as appropriate. Disease Specific survival was analyzed by the Kaplan–Meier method and differences were analyzed by the log-rank test. Statistical significance was displayed in figures at \**p*L<L0.05, \*\**p*L<L0.01, \*\*\**p*L<L0.001, with adjustments for multiple comparisons as appropriate. *P* valueL<L0.05 were considered statistically significant. All statistical tests were two-sided.

## RESULTS

### PNCK Multi-Omics Analysis

A comprehensive and integrative analysis of the cancer genome atlas (TCGA) was performed to identify understudied kinases that were differentially expressed in RCC versus normal (non-cancer) tissues as described in our previous work [7]. Results from the analysis demonstrated that several calcium-calmodulin dependent kinases (CAMK) were overexpressed in RCC tumors (**Supplemental Figure 1A**). Among CAMK members, Pregnancy-Upregulated Non-ubiquitous Calcium/Calmodulin dependent kinase (PNCK or CAMK1B) was found to be the most differentially overexpressed kinase in clear cell renal cell carcinoma (KIRC), with a difference of 6log2-fold between median expression of tumor samples and adjacent normal kidney tissue (**Figure 1A**). We have previously demonstrated that a broader examination of the RNAseq data found PNCK to be in the top 1% of all differentially overexpressed genes in clear cell renal carcinoma [7]. Additionally, PNCK was shown to be significantly overexpressed in papillary (KIRP) (**Figure 1B**) and chromophobe RCCs (**Supplemental Figure 1D**). Other tumors that significantly overexpressed PNCK include lung squamous carcinoma (LUSC), breast carcinoma (BRCA) and hepatocellular carcinoma (LIHC) **(Supplemental Figure 1B-C**). Merging all kidney cancer cohorts, including papillary, clear cell and chromophobe tumors, to a combined kidney cancer dataset (KIPAN), PNCK was shown to be significantly overexpressed. PNCK expression in all cohorts correlated significantly with AJCC stage (**Figure 1D-F**). Kaplan-Meier survival analyses were performed to determine if mRNA levels of PNCK correlated with disease specific survival (**DSS**) (**See Methods**). In Clear Cell (KIRC), Papillary (KIRP) and KIPAN RCC cohorts, elevated PNCK expression was significantly associated with worse DSS (**Figure 1G-I**). TCGA data analyses also revealed a significant correlation between PNCK mRNA levels and other clinical and pathological outcomes, including clinical T staging and histologic grade (Fuhrman Grade) in ccRCC cohorts [15, 20] (**Supplemental Figure 1 E-F)**. PNCK expression quantified via immunohistochemistry of human kidney tumor samples has been correlated to negative survival outcomes in one small study (n=248) [9]. Overall, our analysis is in congruence with previous data that demonstrating PNCK expression in RCC is linked to disease progression and decreased survival.

**Figure 1:**
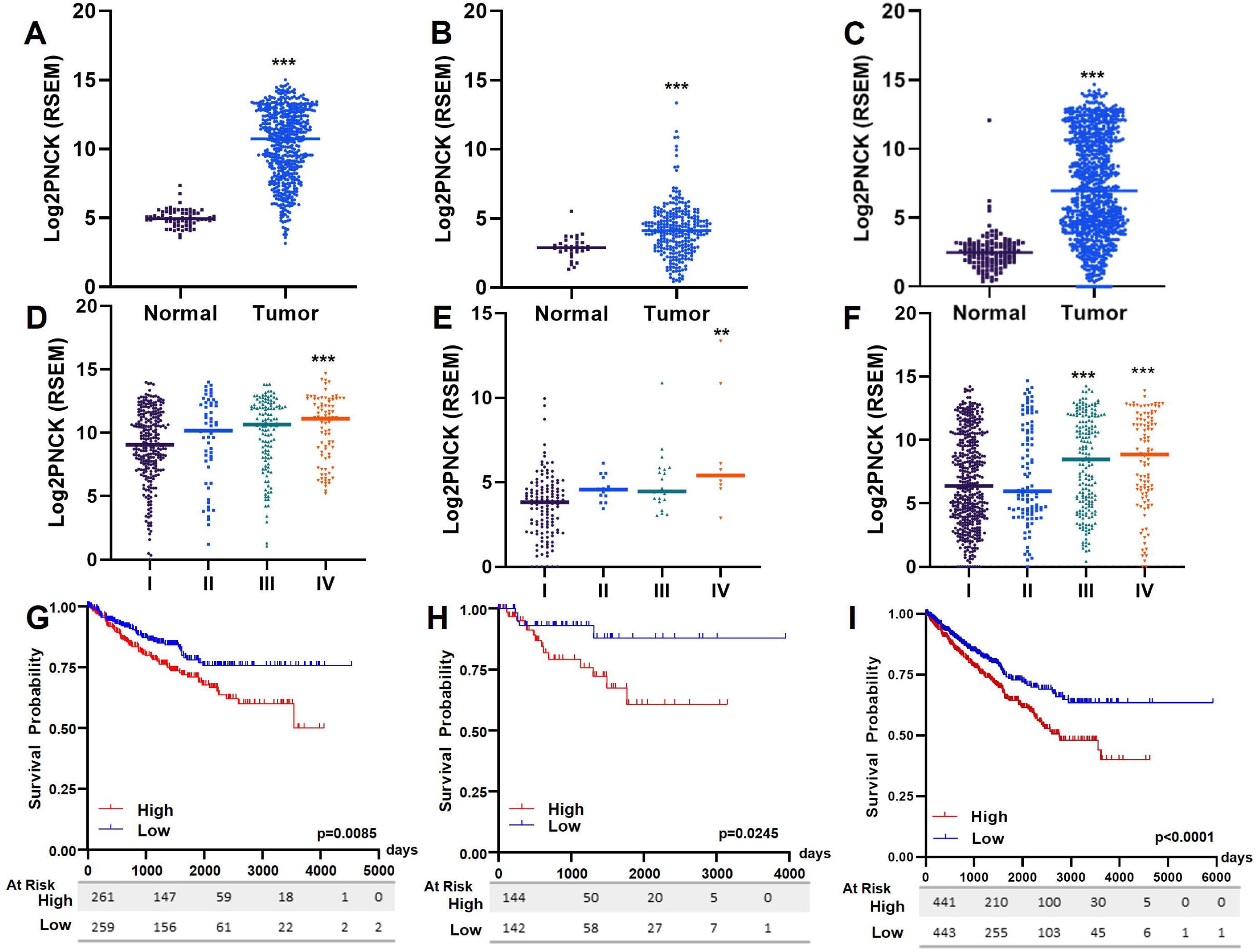
Clinical Relevance of PNCK in Renal Cell Carcinoma From TCGA: (A-C) Normalized expression of PNCK mRNA in Clear Cell (KIRC) (A), Papillary (KIRP) (B) and Pan-Kidney (KIPAN) (C) cohorts. The bar in dot plots represents the median expression. (D-F) Correlation of PNCK expression with clinical and pathological staging in clear cell (D), papillary (E) and pan-kidney (F) RCC cohorts. (G-I). Correlation of PNCK mRNA and Disease Specific Survival (DSS) in Clear cell (G), Papillary (H) and pan-kidney (I) cohorts. (** p<.001, *** p<.0001)

### PNCK Overexpression Leads to Increased levels of phosphorylated CREB with nuclear translocation

PNCK mRNA expression was quantified in a panel of kidney cancer cell lines and 3 control cell lines (HEK293, HUVEC and HREC). Consistent with data from the Cancer Cell Line Encyclopedia (CCLE) [21], PNCK is significantly overexpressed in kidney cancer cell lines compared to normal human renal epithelial cells (HREC) (**Supplemental Figure 2A**). To evaluate the effects of PNCK overexpression to the scale observed in human tumor samples on *in vitro* biological functions, renal cell carcinoma (786-O, A498, ACHN & Caki-1) cells were engineered to stably express PNCK (**See Methods**). Overexpression (OE) was confirmed via qPCR, western blot and immunofluorescence (**Figure 2**). PNCK, unlike other isoforms of CAMK1, lacks the conserved nuclear exclusion signal (NES)[22]. Therefore, we hypothesized that PNCK would be localized to both nuclear and cytoplasmic compartments. Indeed, IF signal of PNCK was intensely nuclear in ACHN and 786-O cells while in A498 and Caki-1 cells, the signal was diffusely cytoplasmic and intensely peri-nuclear. Previous studies have shown that calcium calmodulin dependent kinases (CAMK1 and CAMK2) phosphorylate and activate the transcription factor (Cyclic AMP Response element binding protein) CREB at Ser133 *in vitro*. However, this has never been shown for PNCK[23–26]. We observed that PNCK overexpression led to increased pCREB signal, as measured by immunofluorescence (**Figure 2C, F, I, L)**). While pCREB signal was present both in the control and PNCK OE cells, a more prominent nuclear localization of pCREB was observed in PNCK OE cells while control cells displayed a diffuse (cytoplasmic) signal. CREB localization to the cytoplasm, mitochondria or nucleus is determined by many factors such as pH, hypoxia, and other post-translational modifications including phosphorylation [25, 26]. Signal intensity in the nucleus in PNCK overexpression suggests PNCK may alter sub-cellular location of CREB and the expression of downstream CREB effector genes.

**Figure 2:**
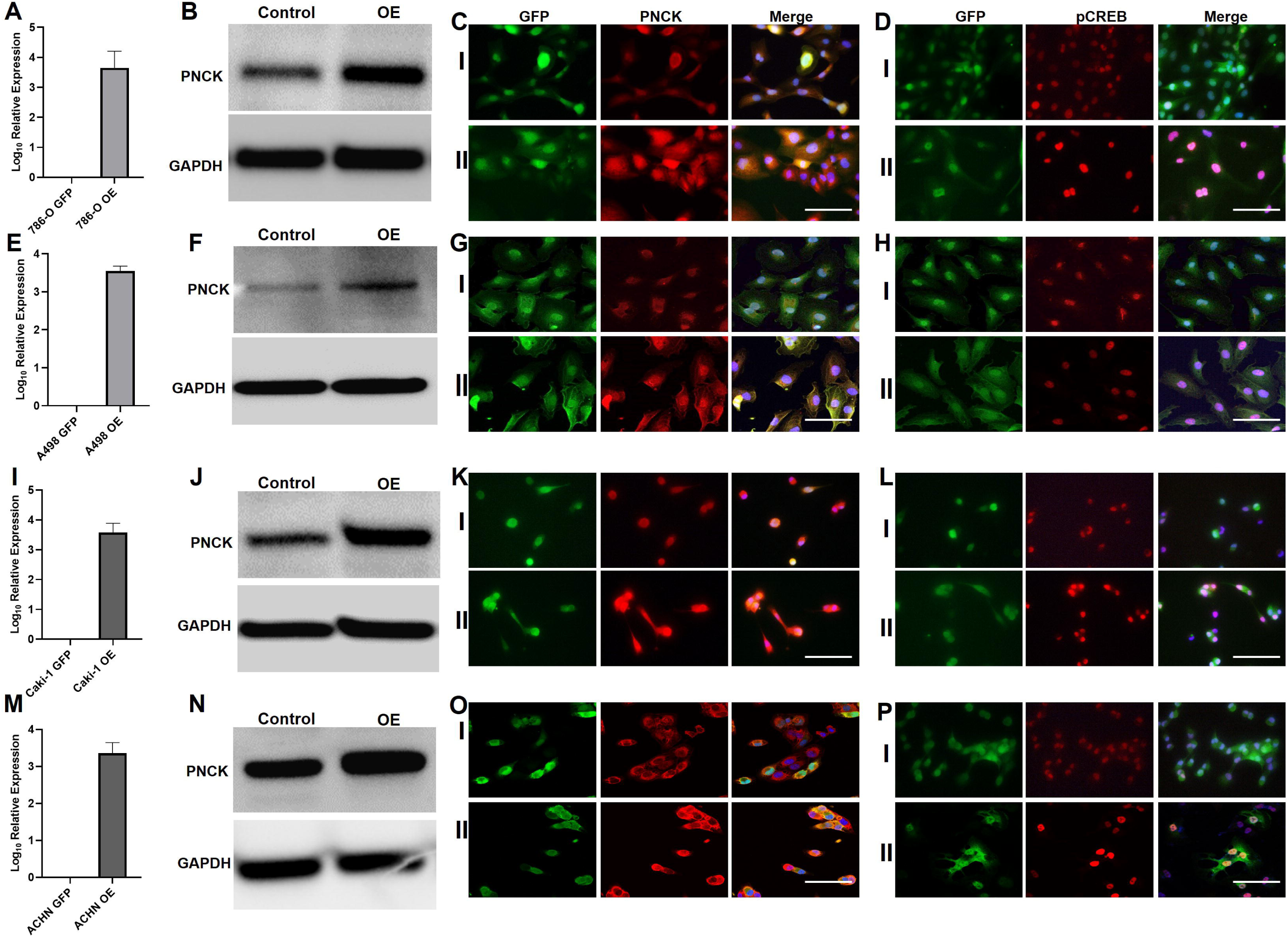
Characterization of PNCK Overexpressing RCC Cell Lines: [A-D 786-O, E-H A498, I-L ACHN, M-P Caki-1] PNCK was stably overexpressed in 786-O (A-D), A498 (E-H), ACHN (I-L), and Caki-1 (M-P) as described in methods. PNCK Overexpression was confirmed by qPCR (A, E, I, M), western blot (B, F, J, N), and Immunofluorescence (C, G, K, O). Changes in Phospho-CREB expression were analyzed by immunofluorescence (D, H, L, P). I: GFP Control Cells, II: PNCK Overexpressing cells. Scale bar represents 10μm.

### Effects of PNCK overexpression on RCC Proliferation and Cell Cycle Progression

To assess the effects of PNCK overexpression on RCC cell growth *in vitro*, rates of cellular proliferation in PNCK OE RCC cells were examined using the real-time xCelligence assay[27] for up to 120 hours (**See Methods**). As predicted, PNCK overexpression led to significant increases in cell growth in all 4 RCC cell lines tested, compared to GFP-transfected controls (**Figure 3A-D**). Notably, PNCK overexpression in HEK-293 cells led to no appreciable difference in proliferation rate or cell index compared to control (**Supplemental Figure 3A**). No appreciable morphological differences were noted in PNCK overexpression cells versus GFP controls thus differences in cell indices and electrical impedance were likely due to differences in cell count. Additionally, growth curves displayed a characteristic “right” shift, with OE cells displaying rapid acceleration of growth in the first 24-48 hours of seeding and reach confluence (As quantified by normalized cell index) faster than GFP controls. After 120 hours, the cell indices for both OE and GFP control generally reach the same height and begin to decrease as cells reach confluence and detach from the sensor (**Data not shown**). Of note, PNCK overexpression also did not have any significant effects on migration or invasion in vitro (data not shown).

**Figure 3:**
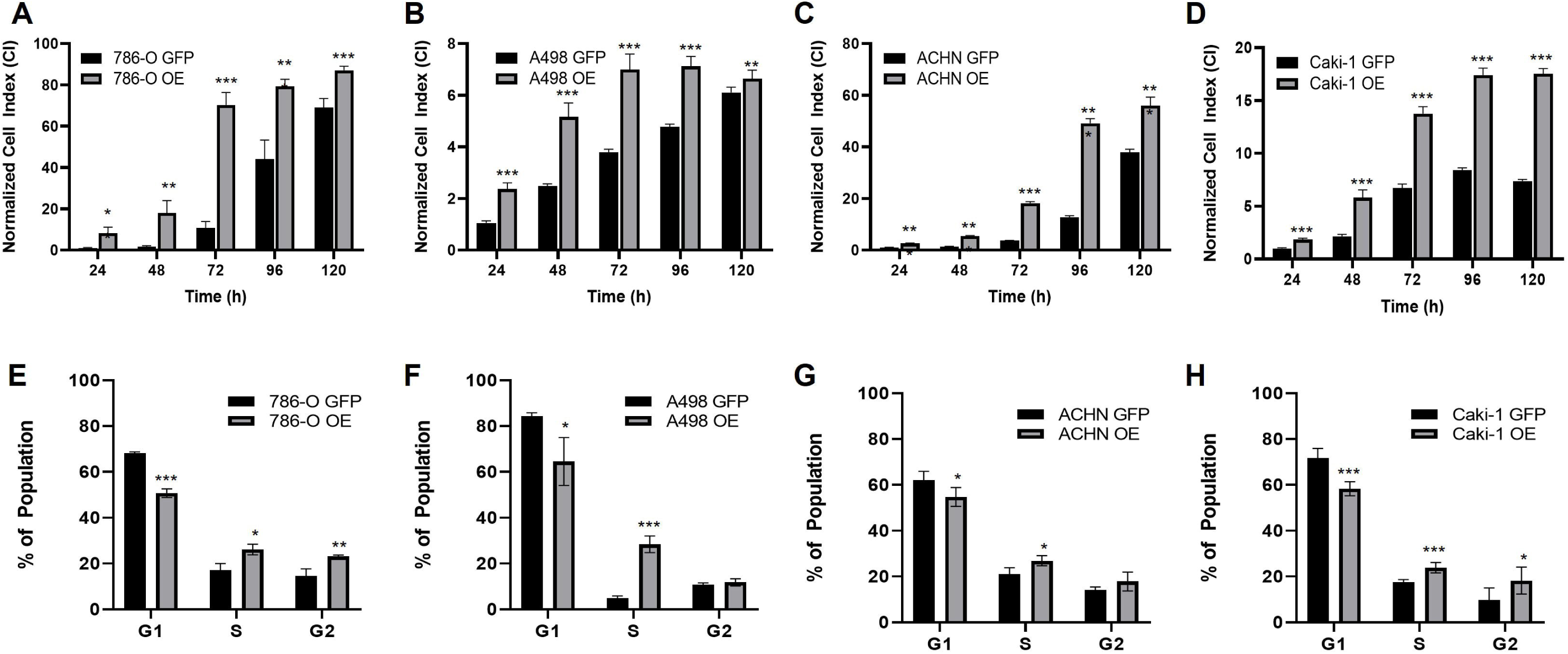
Effects of PNCK Overexpression on RCC Cell Proliferation and Cell Cycle Progression: (A-D) in vitro cell growth of PNCK OE cells and GFP control cells was determined via xCelligence real-time analysis (up to 120h) as described in methods (n=6 per cell line). Cell index was normalized to that of GFP-transfected control cells at each time point. Data is displayed as bar graphs of the average cell index at each time point with the error bars displayed as standard deviation. Experiments were performed in sextuplicate and repeated at least 3 times. *p<.05. **p<.005, ***p<.0001. (E-H) Flow cytometry analysis of PNCK overexpressing and GFP control cells was performed as in methods. Data is displayed as bar graphs of the average percent population in each cell cycle phase with error bars displayed as standard deviation. Experiments were performed in triplicate and repeated 3 times. *p<.05. **p<.005, ***p<.0001.

To further explore the role of PNCK expression in cellular proliferation, flow cytometry was utilized to quantify the population of cells in each stage of the cell cycle (**See Methods**). Previous studies have shown that CAMK1 and CAMK2 partly control G1/S and S/G2 transitions through various mechanisms including activation of CDK/Cyclin complexes[28–32]. In cells that were engineered to overexpress PNCK, there was a significantly higher population of cells in the S and G2 phases and a significantly lower number of cells in the G1 phase compared to GFP controls in 786-O, A498, ACHN and Caki-1 cells (**Figure 3D-F).** In congruence with our proliferation studies, HEK293 cells overexpressing PNCK did not show any changes in cell cycle populations compared to GFP controls (**Supplemental Figure 3B)**. Collectively, these data suggest that PNCK activity likely increases cellular proliferation through effects exerted on cell cycle progression at either G1/S or S/G2 transition.

### Effects of PNCK down-regulation on RCC CREB phosphorylation, Proliferation and Cell Cycle

As overexpression of PNCK led to significant increases in cellular proliferation via progression through cell cycle checkpoints, likely through upregulation of downstream CREB target oncogenes, we hypothesized that knockdown of PNCK would lead to growth arrest. Targeted PNCK inhibition was achieved using double-stranded siRNA (dsiRNA) technology (**See Methods**). RCC cells (786-O, A498, ACHN, Caki-1) were transfected with dsiRNA targeted against the canonical PNCK sequence or Cy3-labeled scramble control. PNCK knockdown was confirmed via qPCR, western blot and immunofluorescence, (**Figure 4**). No significant morphological changes were observed between control and PNCK k/d cells. Compared to Cy3 control, PNCK dsiRNA treated cells had markedly diminished pCREB signal in all RCC cell lines (**Figure 4**).

**Figure 4:**
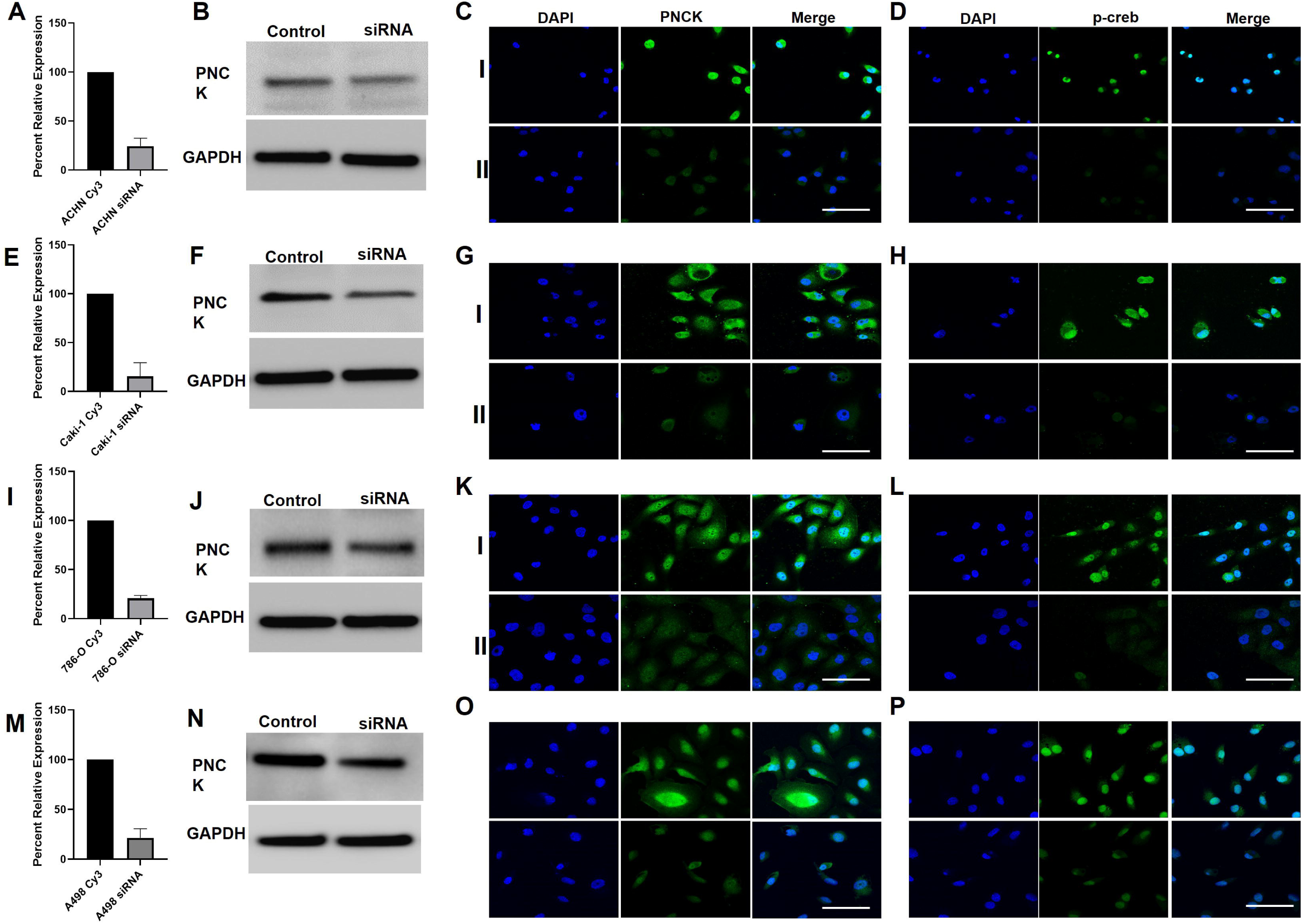
Characterization of PNCK k/d RCC Cell Lines. [A-D 786-O, E-H A498, I-L ACHN, M-P Caki-1] PNCK was transiently knocked down in 786-O (A-D), A498 (E-H), ACHN (I-L), and Caki-1 (M-P) as described in methods. PNCK k/d was confirmed by qPCR (A, E, I, M), western blot (B, F, J, N), and Immunofluorescence (C, G, K, O). Changes in Phospho-CREB expression were analyzed by immunofluorescence (D, H, L, P). I: GFP Control Cells, II: PNCK Overexpressing cells. Scale bar represents 10μm.

Knockdown (k/d) cells were then analyzed using the real time xCelligence proliferation assay. Knockdown cells, compared to dsiRNA scramble control, led to statistically significant inhibition of cell growth *in vitro* (**Figure 5A-D**). The degree and dynamics of growth inhibition differed among cell lines, with ACHN and Caki-1 showing a marked inhibition in proliferation at all time points. Conversely, 786- O cells showed a decrease in proliferation at earlier time points (before 72 hours post transfection), and A498 PNCK k/d was associated with a delayed inhibition in proliferation (beyond 72 hours post transfection) with a noted decrease in normalized cell index between 72-120 hours, suggestive of increased cell death. PNCK k/d did not significantly interfere with RCC migration or invasion in vitro (Data not shown).

**Figure 5:**
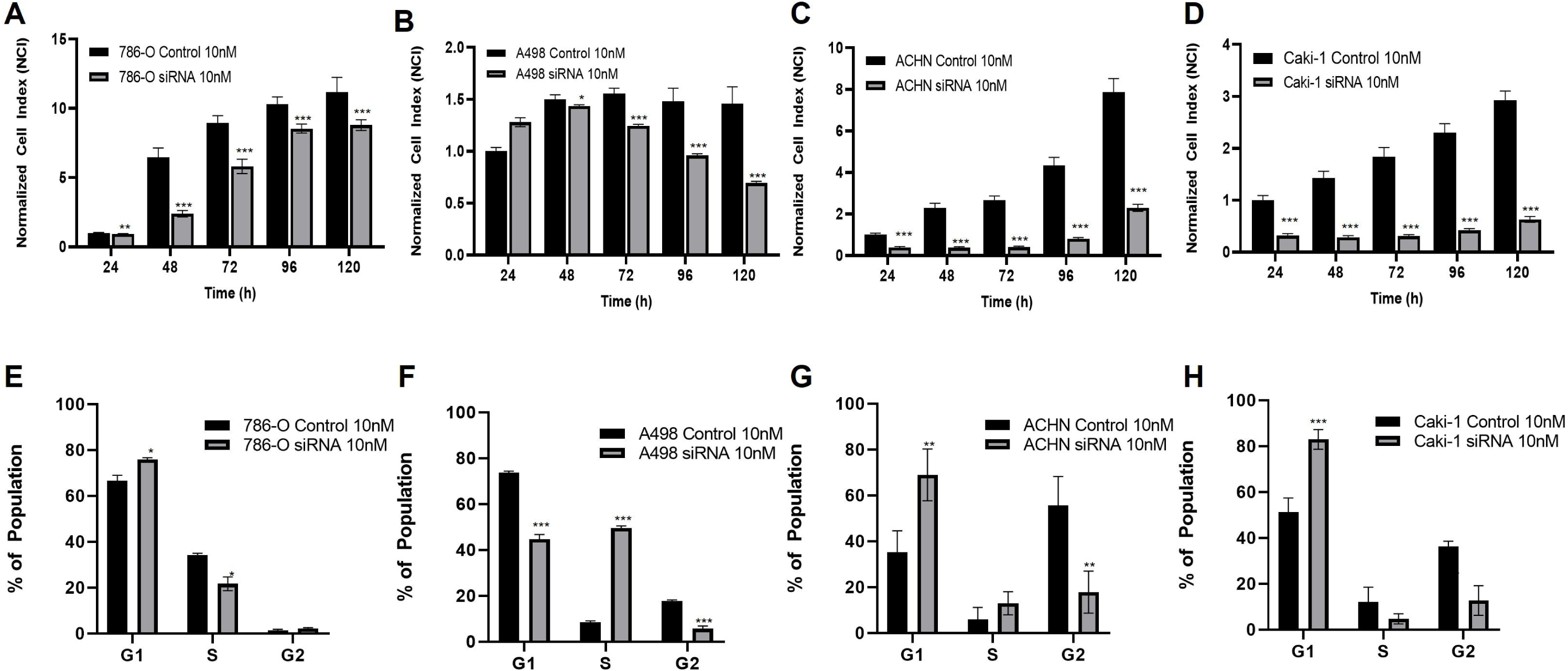
Effects of PNCK Knockdown on RCC Cell Proliferation and Cell Cycle Progression : (A-D) in vitro cell growth of PNCK k/d cells and Cy3 control cells was determined via xCelligence real-time analysis (up to 120h) as described in methods (n=6 per cell line). Cellular index was normalized to that of control cells at each time point. Data is displayed as bar graphs of the average normalized cell index at each time point with the error bars displayed as standard deviation. Experiments were performed in sextuplicate and repeated at least 3 times. *p<.05. **p<.005, ***p<.0001. (E-H) Flow cytometry analysis of PNCK k/d and Cy3 control cells was performed as in methods. Data is displayed as bar graphs of the average percent population in each cell cycle phase with error bars displayed as standard deviation. Experiments were performed in triplicate and repeated 3 times. *p<.05. **p<.005, ***p<.0001.

Next, the effects of PNCK k/d on cell cycle progression were assessed. PNCK knockdown in 786-O, ACHN and Caki-1 cells was associated with G1 cell cycle arrest (with a large population of cells accumulating in G1 and smaller populations in the proliferative S phase and G2 phases) compared to scramble treated controls (**Figure 5E-H**). In A498 cells, cell cycle arrest appeared to occur at the S/G2 transition, with a significant accumulation of cells in the S phase with a very small population observed in G2. Knockdown of PNCK in HEK293 cells was not associated with significant changes in cellular proliferation or cell cycle population (Supplemental Figure 3C,D). To glean insight to the mechanisms of cell cycle arrest, signals of key cell cycle control proteins were analyzed via immunofluorescence. IF studies determined that all k/d cells had increased p21 and p27 signal with decreased cyclin D1and ki67 (**Figure 6**). Differential effects were observed in 786-O cells with decreased cyclin B1 and CDK4/CDK6 signals while no appreciable differences were noted in A498, ACHN or Caki-1 cells (**Supplemental Figure 4**). These data support the hypothesis that PNCK plays a significant role in cell cycle progression at various checkpoints and that inhibition of PNCK and CAMK signaling in kidney cancer cells is cytostatic (as opposed to cytotoxic) via cell cycle arrest.

**Figure 6:**
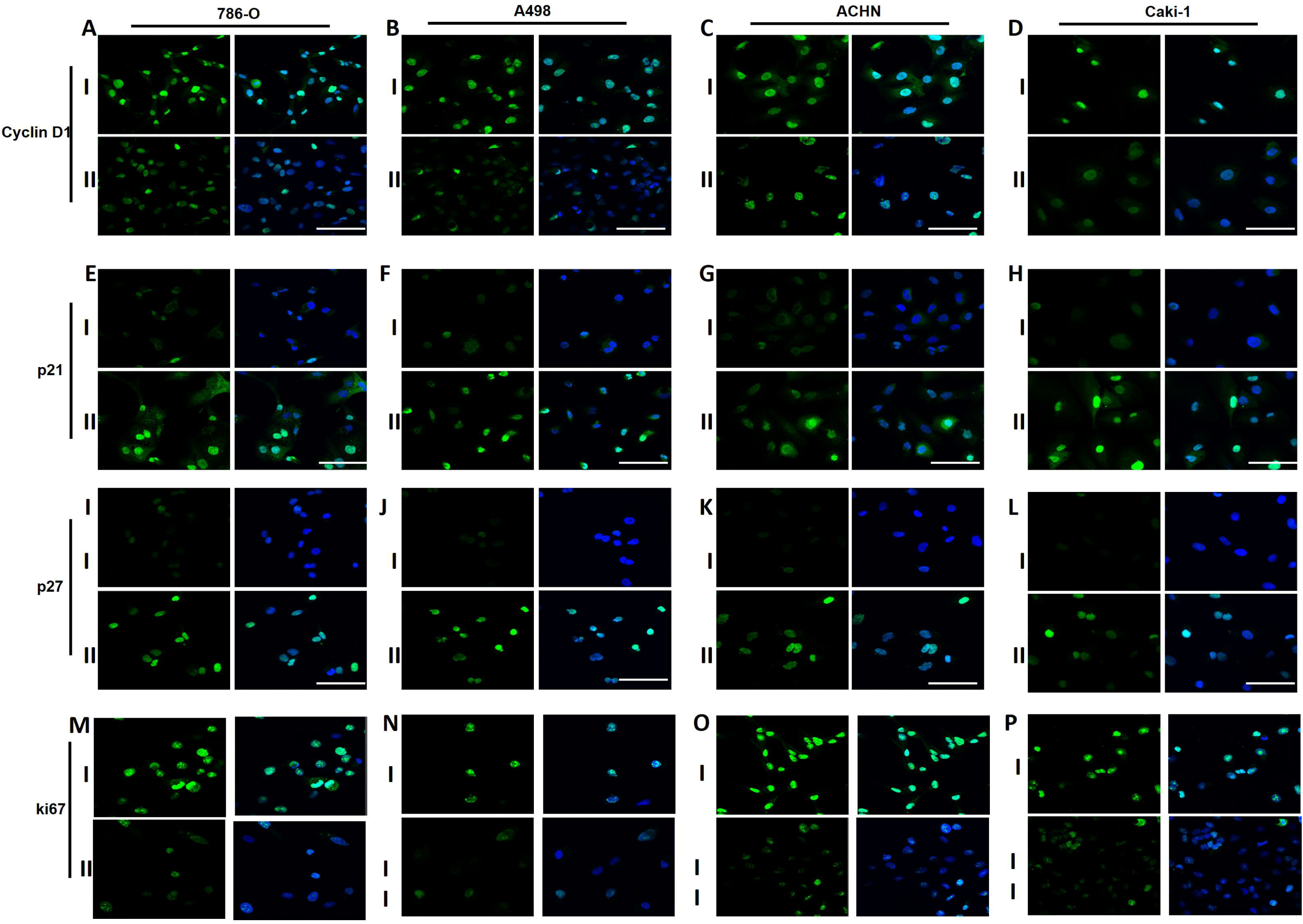
PNCK Knockdown Leads to Changes in Cell Cycle Protein Expression: (A-D) Cyclin D1, (E-H) p21, (I-L) p27, (M-P) ki67. I: Cy3 Control, II: PNCK K/d. (Left panel= Primary antibody, right panel= DAPI merged image). PNCK was knocked down in all 4 RCC Cell lines as described in methods. IF images were taken at 72h post-dsiRNA transfection. Scale bar represents 10μm.

### Knockdown of PNCK in RCC Cells Causes Significant Perturbation of Angiogenesis, Apoptosis, Cell cycle, DNA-Damage response related pathways

To gain a further understanding of the biological functions of PNCK and the pathways through which it exerts its effects, a targeted cancer-pathway gene array analysis was conducted (**See Methods, Supplementary Table 1**). The expression of 84 diverse cancer-related genes representing effector genes in key pathways was assessed upon knockdown of PNCK with dsiRNA.

First, PNCK knockdown was associated with significant changes in the expression of angiogenesis and HIF-1α related pathway genes across all cell lines tested (**Figure 7A)**. Specifically, angiopoietin-1 and angiopoietin-2 were downregulated with the largest magnitudes in each cell line in addition to decreases in FLT1 (VEGFR1), TEK1 (Angiopoietin-1 Receptor), MAP2K1 (MEK1) and PGF (Placental growth factor). PNCK k/d significantly decreased expression of FGF2 and CCL2 in the majority (3 out of 4 cell lines), while expression of VEGFC and KDR were increased in 3/4 cell lines. ARNT (HIF2-B) was only significantly downregulated in VHL wild-type cells, Caki-1 and ACHN. The effects of PNCK k/d on ANGPT1 and ANGPT2 expression were validated at the protein level (by immunofluorescence), showing marked decrease in angipoietin-1 and angiopoietin-2 expression in PNCK k/d vs control cells (**Figure 8**).

**Figure 7:**
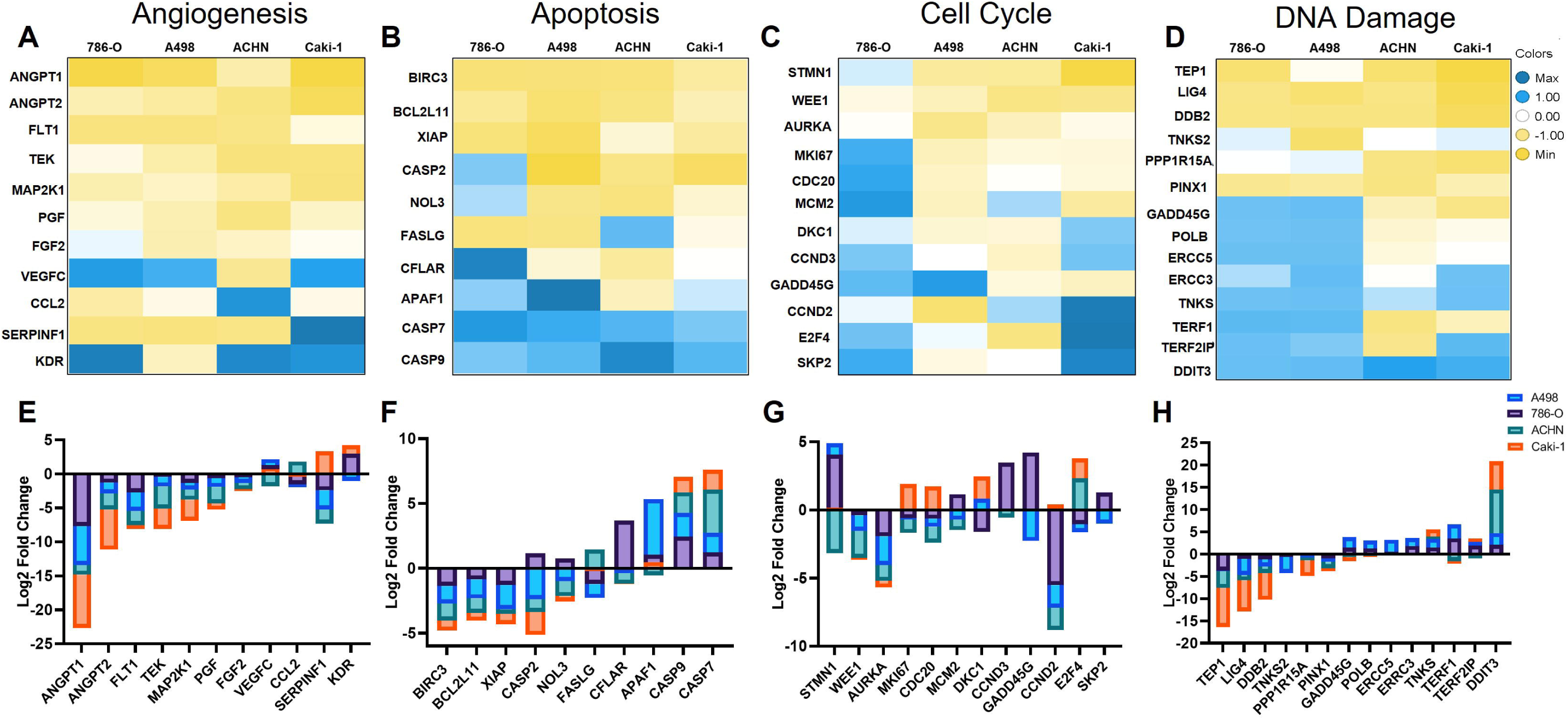
Transcriptional Perturbation of Cancer Pathways Upon PNCK Knockdown: Cy3**-**Control and PNCK knockdown RCC cell lines were generated as in methods. At 48h post-transfection, RNA was extracted from cells and was used to assay 84-cancer pathway related genes in a qPCR array (See Supplemental Table 1) (A-D) Heat Map of differentially expressed Angiogenesis (A), Apoptosis (B), Cell cycle (C) and DNA Damage (D) pathway genes in 4 RCC cell lines. (E-H) Bar plot representation of differentially expressed Angiogenesis (A), Apoptosis (B), Cell cycle (C) and DNA Damage (D) pathway genes in 4 RCC cell lines.

**Figure 8:**
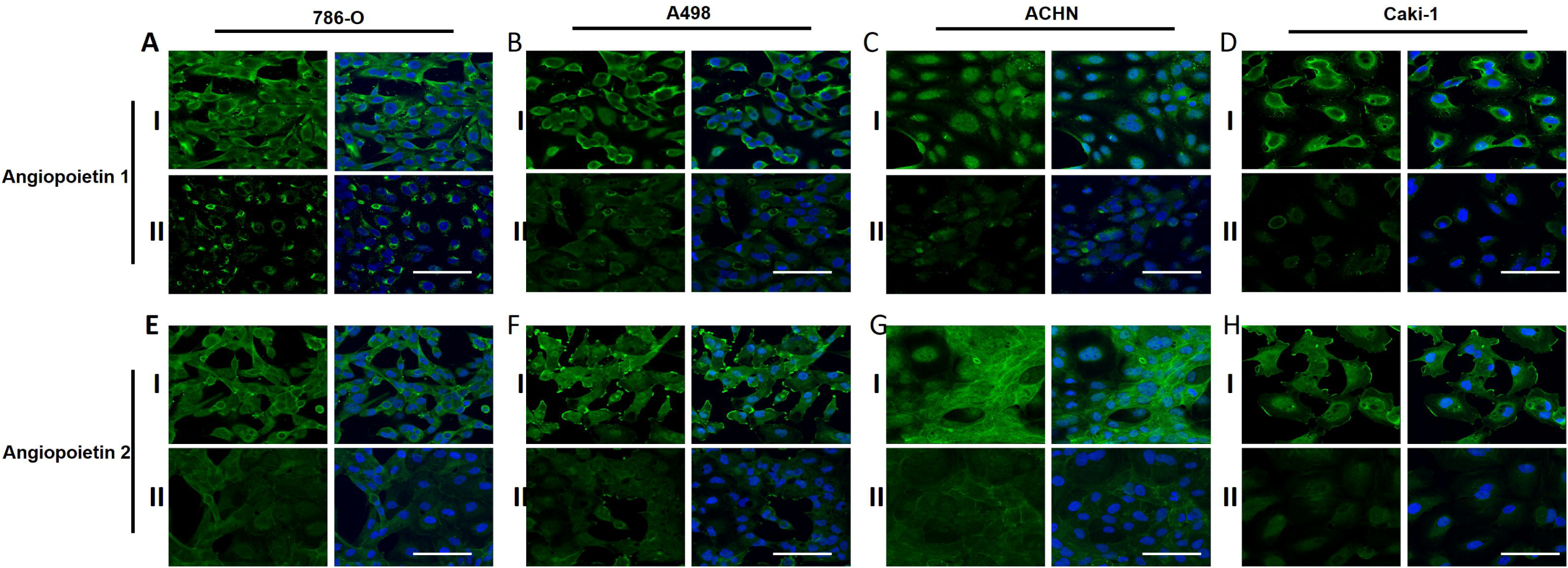
PNCK Knockdown Significantly Alters Angiogenesis Pathways in all 3 Kidney Cancer Cell lines. (A-D) Angiopoietin 1, (E-H) Angiopoietin 2 I: Cy3 Control, II: PNCK K/d. (Left panel= Primary antibody, right panel= DAPI merged image). PNCK was knocked down in all 4 RCC Cell lines as described in methods. IF images were taken at 72h post-dsiRNA transfection. Scale bar represents 10μm.

Second, PNCK knockdown resulted in significant modulation in apoptosis related genes, independent of VHL mutational status (**Figure 7B**). For example, PNCK knockdown led to significantly increased expression of the proapoptotic genes CASP7 and CASP9 in all cell lines and increase in the pro-apoptotic APAF-1in 3 out of 4 cell lines. Significant down regulation in the expression of the anti-apoptotic genes BIRC3, BCL2L11 and XIAP were observed in all cell lines, while decrease in the anti-apoptotic NOL-3 was observed in 3 of 4 cell lines. The above gene expression changes were associated with increased RCC apoptosis, as demonstrated by significant increase in caspase 3 and 7 activity *in vitro* in 786-0, A498, ACHN and Caki-1 upon PNCK down regulation (**Figure 9**).

**Figure 9:**
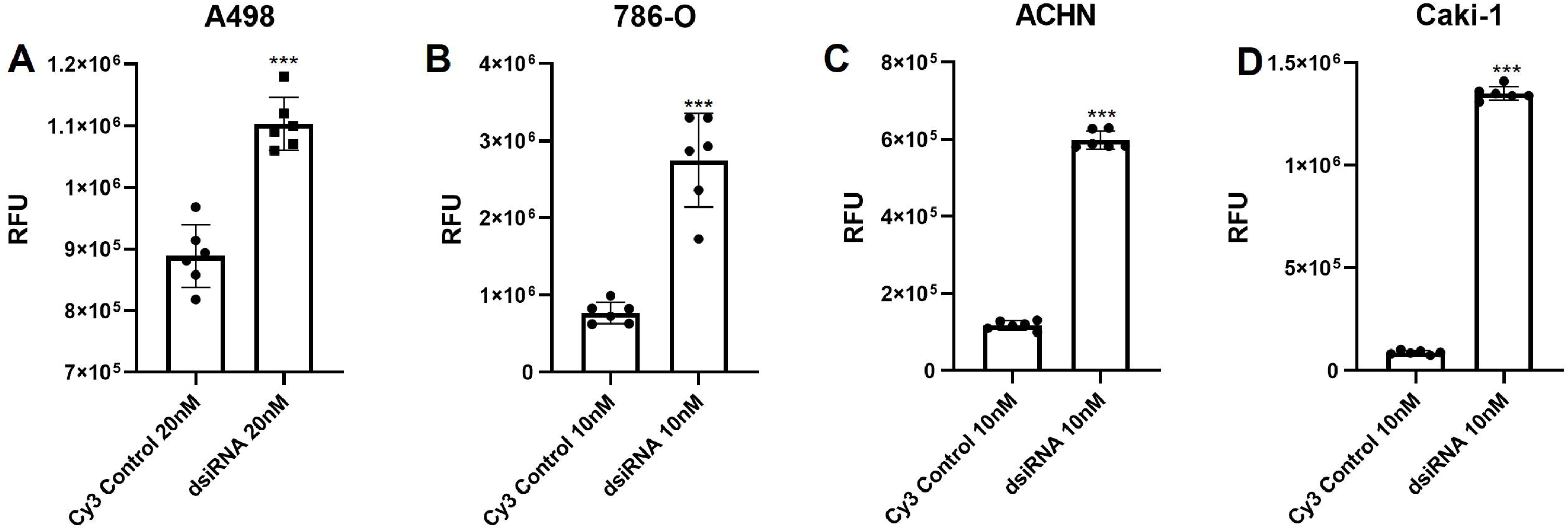
Effects of PNCK Knockdown on Caspase Activity: RCC cells treated with either dsIRNA targeting PNCK or a Cy3-labeled scrambled control were assayed at 72h post-transfection for caspase 3/7 activity using Promega Caspase Glo 3/7 Apoptosis Assay (see Methods). Luminescence was detected by Promega luminometer and displayed as RFU. Data displayed as bar-dot plot +/- standard deviation. (n=6). *p<.05. **p<.005, ***p<.0001.

Third, in congruence with previous studies linking PNCK to the DNA damage-response, we found that PNCK k/d led to significant perturbations in DNA damage response and chromosomal and teleomeric integrity pathway genes (**Figure 6 C**). Across all cell lines tested, TEP1, LIG4 and DDB2 and PINX1 were all significantly downregulated, with Caki-1 showing >-440 fold difference in expression of DDB2. Conversely, all cell lines showed significant overexpression of DDIT3 (DNA Damage Inducible Transcript 3) also known as CHOP, as well as TNK5.

Cell-cycle related genes demonstrated variability across the 4-cell lines tested with some consensus around significant downregulation of WEE1[33] and AURKA[34], involved in cell cycle regulation (**Figure 6D**). Expression of MKI67, CDC20 and STMN1 were significantly decreased in 3 of 4 cell lines upon PNCK k/d. 786-O, a cell line with p53 mutation, was a clear outlier with upregulation of many cell-cycle related genes in response to PNCK knockdown. Interestingly, common changes in gene expression were observed only in VHL mutant cell lines (786-0 and A498), including significant upregulation of the tumor suppressors GADD456 and TERF1, as well as POLB and ERCC5. Our previously mentioned experiments confirmed k/d of PNCK leads to cell cycle arrest at G1 though the distinct mechanisms are likely different between p53-mutant and p53 wild-type cell lines. However, decreases in cyclin D1 and increase in p21/p27 were consistent across all cell lines (**Figure 6**).

## DISCUSSION

The human genome encodes approximately 634 kinases, of which only 49 are current targets of FDA approved oncological drugs [6]. Renal cell carcinoma (RCC) is an example of a human malignancy where (7 FDA approved) kinase inhibitors have led to improved outcomes, but not cures [35, 36]. During the last decade, kinase inhibitor research in RCC has been limited to developing more potent or selective antiangiogenic TKIs in combination with checkpoint inhibitors [36], while overlooking potentially promising non-TK targets. This status quo has prevented progress in the development of novel kinase inhibitor in RCC. Despite several studies associating Calcium/Calmodulin kinase activity with cancer disease progression, there has never been a small molecule inhibitor in clinical trials that targets this pathway[28, 31]. There are no small molecule inhibitors and no experimentally derived crystal structure for PNCK; and its substrate is not known. Yet, as most kinases, PNCK and other calcium/calmodulin-dependent protein kinases are likely targetable by small molecules and therefore potential drug targets for the development of novel first in class cancer therapeutics.

The current study demonstrates the clinical and biological relevance of PNCK, an understudied kinase, in renal cell carcinoma. Using data from TCGA, PNCK was shown to be the most overexpressed kinase in RCC, at higher levels than well-established kinases currently targeted by FDA approved anti RCC agents[7]. PNCK was significantly overexpressed in both clear cell and importantly, papillary RCC, an RCC subtype lacking highly effective therapies. Not only PNCK was associated with tumor grade and stage, but importantly, its overexpression was associated with poor disease specific survival (**Figure 1**). The above results validate a prior study with smaller number of patients, where overexpression of PNCK (by IHC determination) was found to be an independent negative prognostic factor[9]. Moreover, our observations support other reports demonstrating the prognostic significance of PNCK in other cancers, such as breast, hepatocellular and nasopharyngeal carcinoma[7, 8, 10, 12, 17].

PNCK plays a critical role in RCC proliferation *in vitro*, as demonstrated by promotion of cell growth when PNCK was overexpressed, while growth was significantly inhibited upon PNCK knockdown. Inhibition of cell growth by PNCK knockdown was associated with cell cycle arrest and apoptosis. These effects were observed across the 4 cell lines studied, including VHL mutant (786-0, A498) and VHL wild type (Caki-1) clear cell RCC lines, as well as in the papillary RCC cell line, ACHN, underscoring the biological relevance, and therapeutic potential of PNCK inhibition across different RCC subtypes. Interestingly, the effects of PNCK overexpression and knockdown were absent in control HEK293 cells, suggesting that cancer-promoting PNCK activity may require an oncogenic milieu.

PNCK was found to modulate CREB phosphorylation, as demonstrated by increased pCREB signal (by IF) in PNCK overexpressing, while its levels decreased in PNCK k/d RCC cells. CREB is a transcription factor known to play a critical role in oncogenesis and progression in multiple cancers, including renal cell carcinoma[25, 37, 38].Wang et al reported that targeting CREB is associated with inhibition of RCC growth and metastatic abilities[38]. CREB regulates the expression of several cell cycle checkpoints and is crucial in cell cycle progression in cancer [39–41]. In particular, the expression of both cyclins A and D1, which control the G1/S transition, are highly dependent on CREB activity with cAMP response elements found in the promoters. CREB inhibition was found to induce cell cycle arrest in the S phase, followed by apoptosis in esophageal cancer cells[42, 43], while treatment with a CREB inhibitor in NSCLC caused arrest at the G2/M transition[43]. Our results have shown that PNCK k/d led to decreased pCREB signal in all 4 RCC cell lines. This caused arrest at G1 in 786-O, ACHN and Caki-1 cells, while it caused arrest in the S-phase in A498 cells (**Figure 5**). Previous work in MCF-7 cells has shown that knockdown of CAMK1 led to decreased expression of cyclin D1 and hypophosphorylated rb, with increased expression of cell cycle inhibiting proteins, p21 and p27, causing cell cycle arrest at G1. We have demonstrated here that PNCK (an isoform of CAMK1) k/d resulted in decreased cyclin D1 and increased p21 and p27 expression. The current report provides first time evidence that PNCK inhibition in renal cell carcinoma induces cell cycle arrest via modulation of CREB signaling. Gene expression studies (**Figure 6**) found that PNCK additionally down regulates WEE-1 and AURKA, genes that play a critical role in cell cycle regulation. Further immunofluorescence studies also showed that PNCK k/d resulted in decreased levels if ki67, cyclin A and cyclin B.

PNCK down regulation induced apoptosis in all 4 RCC cell lines, as supported by PNCK k/d mediated significant upregulation of pro-apoptotic genes and down regulation of anti-apoptotic genes (**Figure 7B**). These findings were further validated by demonstration of increased caspase 3 and 7 activity in PNCK k/d 786-0, A498, ACHN and Caki-1 cells (**Figure 9**). It has previously been shown that PNCK knockdown and knockout in nasopharyngeal carcinoma induced apoptosis both *in vitro* and *vivo*[10]. Transcriptomic analyses found significant alterations to the PI3K/Akt/mTOR signaling pathways, offering a clue to the mechanism of apoptosis. Our western blot analyses only saw decreased pAkt signal in VHL-mutant cell lines, A498 and 786-O (Data not shown). The mechanism by which PNCK downregulation causes apoptosis is likely multifactorial, though centered around downregulation of CREB mediated genes, prolonged cell cycle arrest and increased DNA damage burden.

Prior work suggests that PNCK plays a role in the DNA damage pathway, as knockout of PNCK sensitized cells to DNA damaging chemotherapies such as carmustine and temozolomide[44] and also led to increased chromosomal instability[45]. Knockdown of PNCK resulted in loss of a reporter HAC (human artificial chromosome) in yeast cells. PNCK knockdown also caused a significant increase in double stranded breaks and nucleoplasmic bridges-a sensitive measure of chromosome damage. These prior studies support our cancer pathway analyses which show altered expression of genes involved in the DNA damage response, chromosomal and telomeric stability and anti-apoptosis pathways with an increase in expression of pro-apoptotic genes. PNCK knockdown led to significant decreases in DDB2 and LIG4 and in all 4 RCC cell lines. DDB2 encodes for DNA Damage Binding Protein 2 while LIG4 encodes for DNA Ligase 4[46]. These two proteins are both integral for the DNA damage response pathway, inhibit apoptosis, and promotes chemoresistance[46]. DDB2 has been shown to be overexpressed in triple-negative breast cancer cells and knockdown with siRNA led to cell cycle arrest and apoptosis[47], likely due to the accumulation of damaged DNA. LIG4 is involved in double stranded DNA breaks and NHEJ and is frequently overexpressed in tumors[48]. Conversely, the expression of DDIT3 (DNA Damage Inducible Transcript 3), the gene that encodes for CHOP, was significantly increased in all 4 cell lines in response to PNCK knockdown. Overexpression of CHOP has been linked to cell cycle arrest and also induces apoptosis[49]. Therefore, it is highly likely that PNCK activity controls expression and activity of DNA damage response pathways and that knockdown both leads to accumulation of DNA damage which triggers apoptosis, arrests the cell cycle and also likely sensitizes cells to DNA-damaging agents. This is of particular interest in the treatment of RCC and other chemo-and radio-insensitive tumors as pharmacological PNCK inhibition may work to sensitize tumors to DNA damaging agents or other therapies which target the cell cycle via orthogonal pathways.

Renal cell carcinoma, especially clear cell RCC, is a predominantly angiogenic disease [50], and oral antiangiogenic (anti-VEGF) tyrosine kinase inhibitors are the backbone of modern therapies for advanced disease [4]. Therefore, targeting angiogenesis by thwarting the downstream effects of HIF1-α are imperative for treatment. A previous study in breast cancer models discovered a long-noncoding RNA which was shown to activate PNCK which phosphorylates IkBa at Ser32 and triggers calcium-induced NF-kB signaling activity[8]. Knockdown of the lncRNA resulted in decreased angiogenesis in PDX models. HIF1-α gene expression is known to be controlled by NFKB[51]. Therefore, we hypothesize that decreased angiogenesis from PNCK k/d is likely mediated via NFkB and CREB through decreased HIF-1a downstream genes. Indeed, PNCK k/d was associated with marked down regulation of hypoxia and angiogenesis signaling pathways. PNCK k/d significantly decreased expression of HIF1-a target genes such as CA9, ANRT, LDH ANGPT1, ANGPT2, TEK, PGF, MAP2K1 and FLT1 (**Figure 6**). Significant inhibition of Angiopoietin-1 and 2 expression was confirmed in all PNCK k/d RCC lines tested, by immunofluorescence (Figure 8). Conversely, PNCK k/d led to increases in expression of KDR and VEGFC in several cell lines. The prominent effects observed (at the gene expression and protein levels) of PNCK k/d on angiopoietin 1 and 2, reported for the first time in this study, strongly suggest that these two angiogenic cytokines are important targets of PNCK and that PNCK may regulate tumor angiogenesis via angiopoietin signaling. Importantly, our results strongly support the concept of targeting PNCK as an alternative, non-canonical antiangiogenic strategy, especially in the setting of advanced RCC refractory to currently approved anti-VEGF receptor tyrosine kinase inhibitors.

In summary, this is the first report to our knowledge characterizing the cellular and molecular effects of PNCK and PNCK inhibition in renal cell carcinoma. Our results demonstrate that PNCK has both clinical and biological relevance in this fatal disease. We showed that PNCK inhibition exerts direct antitumor effects, by inhibition of cell growth, induction of RCC cell cycle arrest and apoptosis, as well as indirect effects, by regulating expression of angiogenesis and DNA damage response pathways. The above data strongly suggest that PNCK represents a potentially promising target for novel, urgently needed RCC biotherapies, and warrant further research -the current focus of our laboratories-aimed at the development of small molecule PNCK inhibitors as well as in vivo and preclinical studies aimed at targeting PNCK in RCC and other cancers.

## Supplemental Data

**Supplemental Figure 1: TCGA Analysis of PNCK Expression**. A) Expression of Calcium-Calmodulin dependent kinases (CAMKs) in Renal Cell Carcinoma Cohorts. B) Expression of PNCK in Solid Tumors. PNCK is expressed highly in Clear Cell (KIRC), Lung Squamous (LUSC), Prostate (PRAD), Bladder (BLCA), Chromophobe (KICH) and Papillary (KIRP) carcinomas. C) Differential expression of PNCK in solid tumors. PNCK is most differentially overexpressed in Clear cell (KIRC), Lung squamous (LUSC) and Liver Hepatocellular (LIHC) carcinomas. D) PNCK differential expression in Kidney Chromophobe tumors (KICH). PNCK expression in Clear Cell (KIRC) tumors correlates significantly with T-stage progression (E) and Fuhrman Grade (F).

**Supplemental Figure 2: Expression of PNCK mRNA in RCC Cell lines relative to expression in normal kidney endothelial cells (HREC)**

**Supplemental Figure 3: Effects of PNCK Overexpression and Knockdown in HEK293 Cells.** A) xCelligence normalized cell index of HEK293 GFP control cells versus PNCK OE. B) Flow cytometry analysis of cell cycle populations in HEK293 GFP control cells versus PNCK OE. C) xCelligence normalized cell index of HEK293 Cy3 control cells versus PNCK k/d. D) Flow cytometry analysis of cell cycle populations in HEK293 Cy3 control cells versus PNCK k/d

**Supplemental Figure 4: Immunofluorescence Analysis of Cell Cycle-related Proteins in PNCK Knockdown Cells**. (A-D) Cyclin A, (E-H) Cyclin B, (I-L) CDK4, (M-P) CDK6. I: Cy3 Control, II: PNCK K/d. (Left panel= Primary antibody, right panel= DAPI Merged image). PNCK was knockdown in all 4 RCC Cell lines as described in methods. IF images were taken at 72h post-dsiRNA transfection. Scale bar is 10 microns.

**Supplemental Table 1: RT2 Cancer Profiler Array Data**

## ACKNOWLEDGEMENTS

This work was supported by NIH grants U54HL127624 (BD2K LINCS Data Coordination and Integration Center, DCIC), U24TR002278 (Illuminating the Druggable Genome Resource Dissemination and Outreach Center, IDG-RDOC), U01LM012630 (BD2K, Enhancing the efficiency and effectiveness of digital curation for biomedical ‘big data’), and P30CA240139 (NCI Sylvester Cancer Center Support Grant), and the State of Florida Biomedical Research Program, Bankhead Coley grant 9BC13 .

