## Supplementary figures and images for "Cellular And Molecular Effects Of Understudied Kinase Pregnancy-Upregulated Non-Ubiquitous Calcium-Calmodulin Dependent Kinase (PNCK) In Renal Cell Carcinoma"

### Supplemental Figure 1

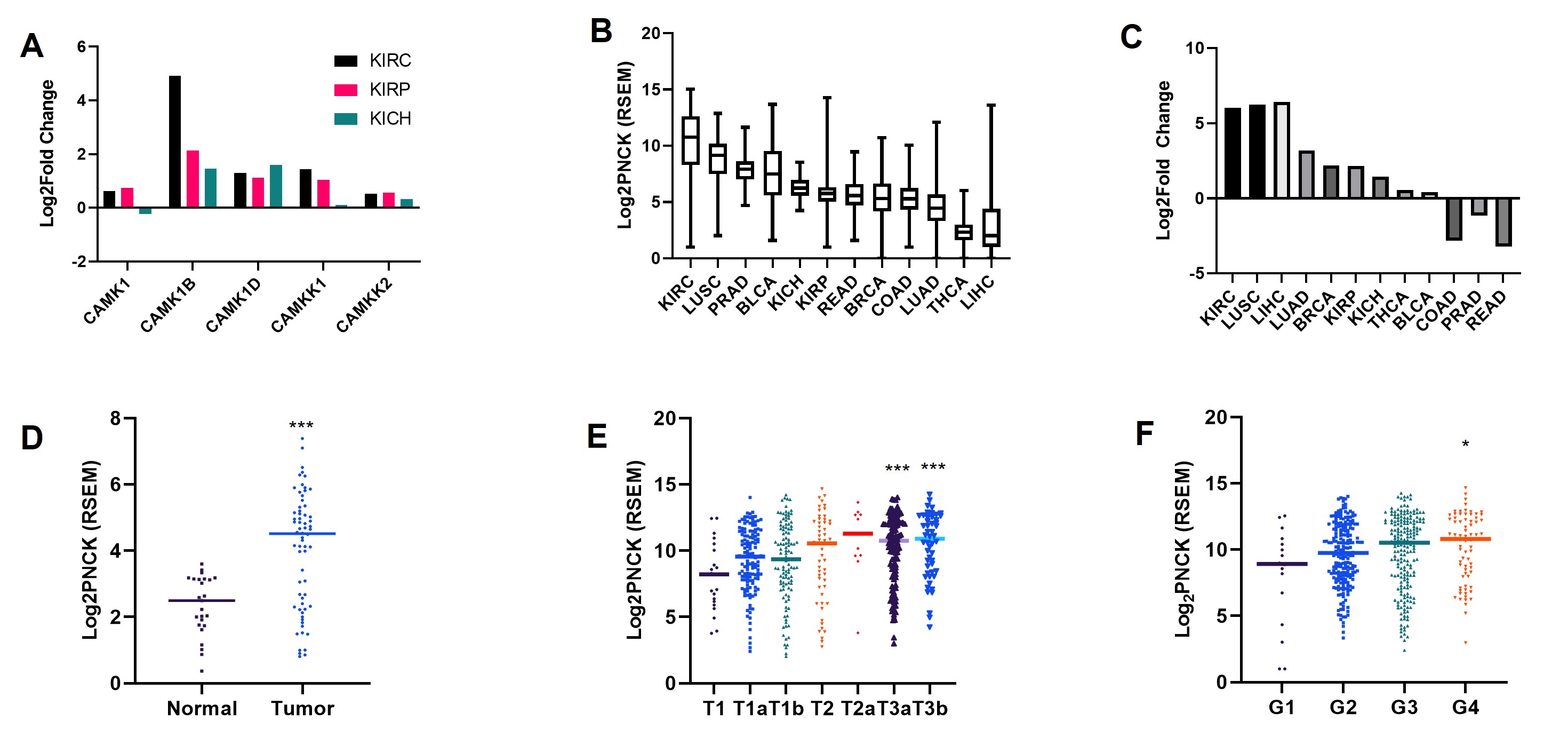

### Supplemental Figure 2

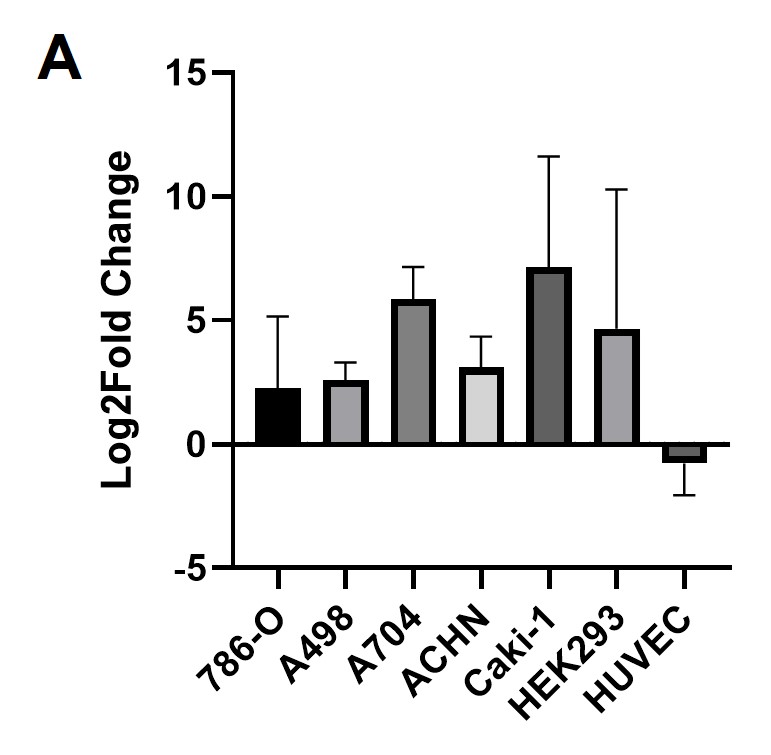

### Supplemental Figure 3

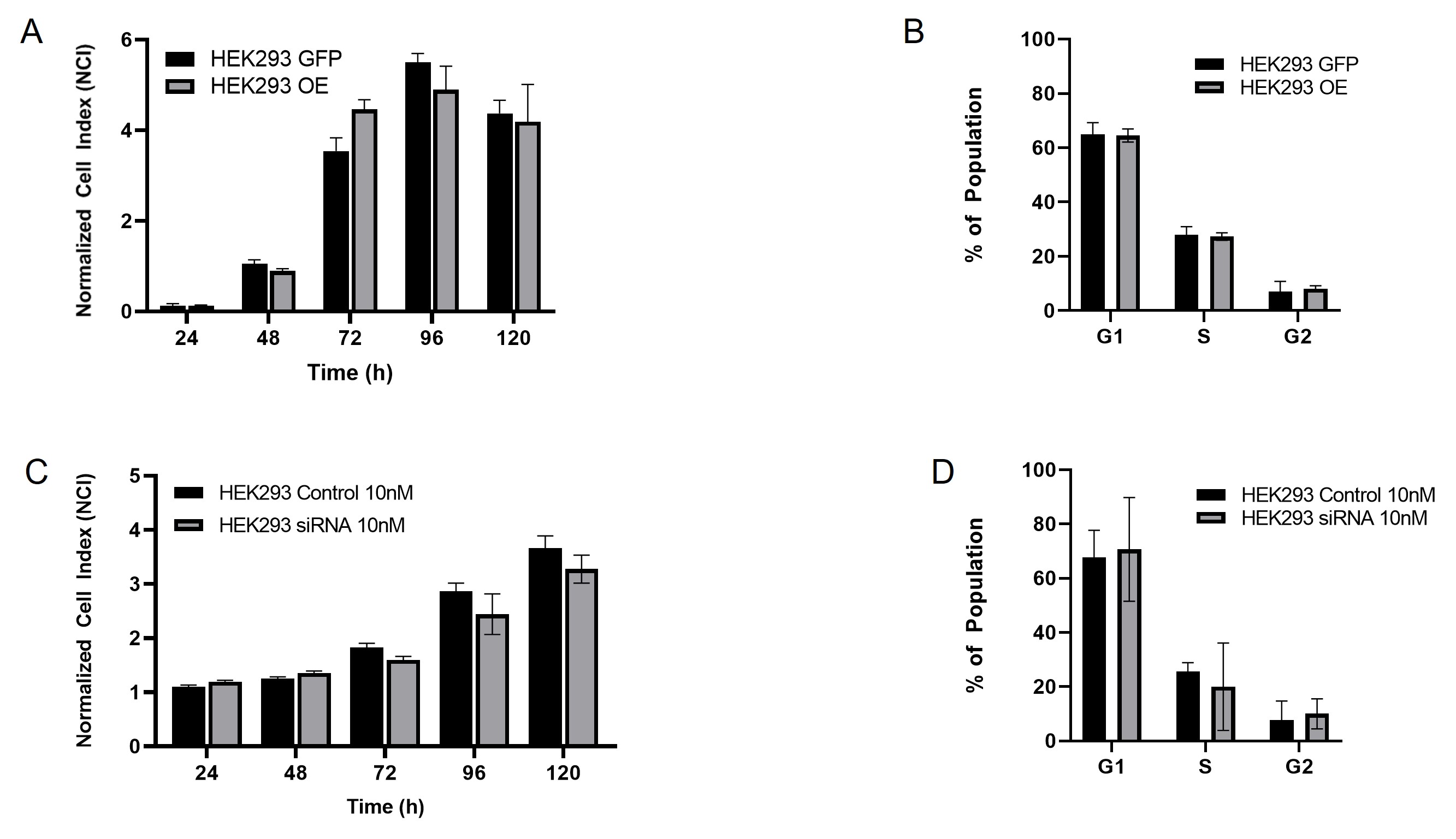

### Supplemental Figure 4

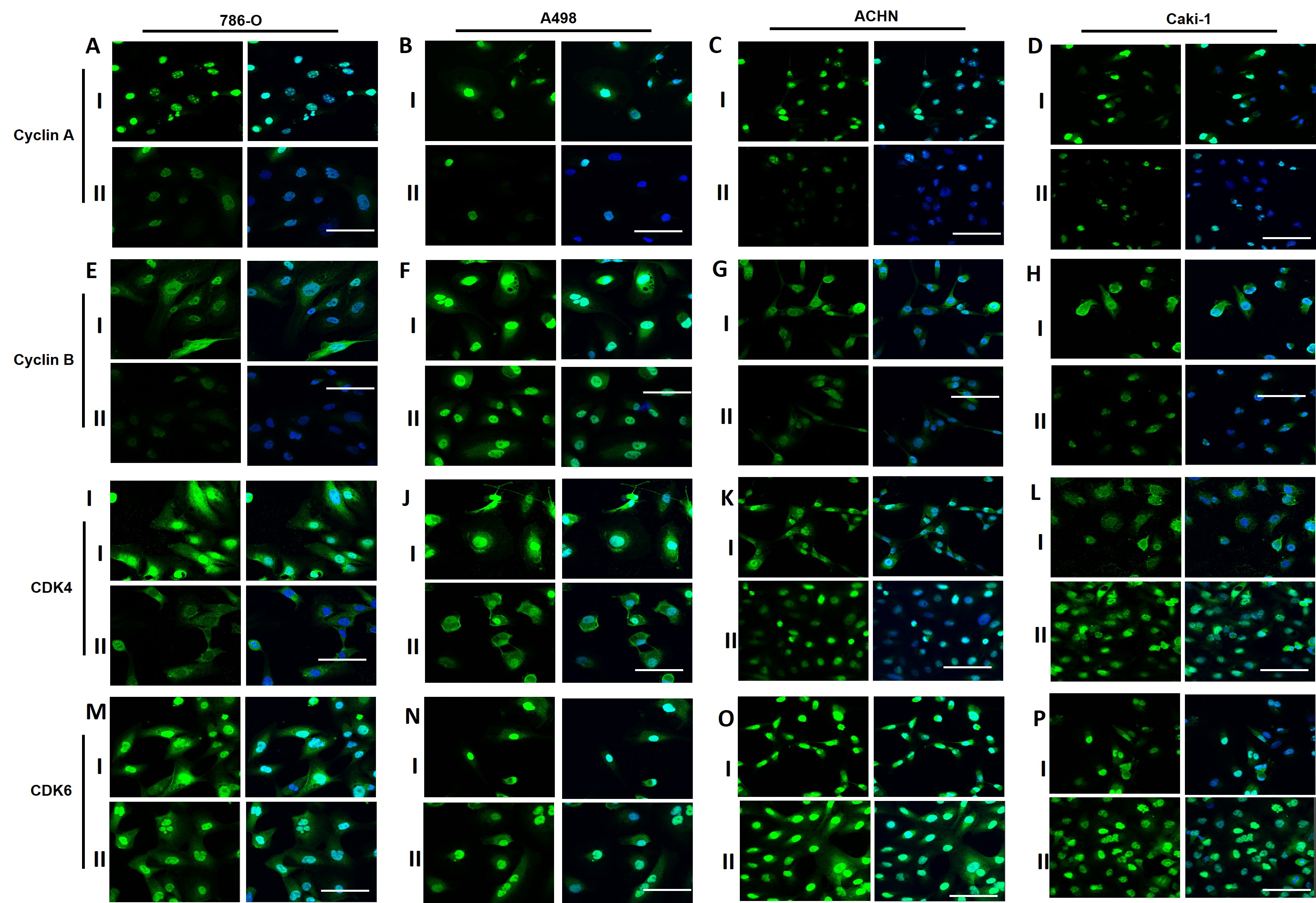
